# A whole genome analysis of the red-crowned crane provides insight into avian longevity

**DOI:** 10.1101/188656

**Authors:** HyeJin Lee, Oksung Chung, Yun Sung Cho, Sungwoong Jho, JeHoon Jun, Jessica A. Weber, Jungeun Kim, Jeongheui Lim, Jeremy S. Edwards, Woon Kee Paek, Jong Bhak

## Abstract

The red-crowned crane (*Grus japonensis*) is an endangered and large-bodied crane native to East Asia. It is a traditional symbol of longevity and its long lifespan has been confirmed both in captivity and in the wild. Lifespan in birds is positively correlated with body size and negatively correlated with metabolic rate; although the genetic mechanisms for the red-crowned crane’s long lifespan have not previously been investigated. Using whole genome sequencing and comparative evolutionary analyses against the grey-crowned crane and other avian genomes, we identified candidate genes that are correlated with longevity. Included among these are positively selected genes with known associations with longevity in metabolism and immunity pathways (*NDUFA5, NDUFA8, NUDT12 IL9R, SOD3, NUDT12, PNLIP, CTH*, and *RPA1*). Our analyses provide genetic evidence for low metabolic rate and longevity, accompanied by possible convergent adaptation signatures among distantly related large and long-lived birds. Finally, we identified low genetic diversity in the red-crowned crane, consistent with its listing as an endangered species, and we hope this genome will provide a useful genetic resource for future conservation studies of this rare and iconic species.

## Introduction

The red-crowned crane (*Grus japonensis*) is one of the rarest cranes in the world and is a symbol of longevity in East Asia [1]. It has a maximum lifespan of 30 years in the wild and 65 years in captivity [2], which is substantially longer than the average lifespan recorded in 144 other avian species (19 and 24 years, respectively) (Supplementary Table S1). It is also one of the largest avian species [3], measuring on average 150-158 cm long with a 220-250 cm wingspan [4], and weighing on average 4.8 to 10.5 kg [5]. Since avian body sizes are known to be positively correlated to lifespan and negatively correlated to metabolic rate [6, 7], it has previously been suggested that the longevity of the red-crowned crane is related to its low metabolic rate [8].

There are currently two primary red-crowned crane populations, including a non-migratory population in northern Japan and a migratory continental group that ranges across southeastern Russia, northeastern China, eastern Mongolia, and Korea [9]. Also a symbol of fidelity in East Asia, the red-crowned crane typically pairs for life and performs elaborate synchronized courtship dances. It relies on wetlands for breeding and the loss and pollution of these habitats has resulted in severe population declines [10]. With a total of only around 3,000 individuals in 2017, coupled with long generational lengths (12 years), it is one of the most endangered avian species [9]. It has been classified as threatened since 2000 and has been classified as ‘endangered’ on the International Union for Conservation of Nature Red List [11]. Conservation genetic studies have primarily focused on microsatellite and mitochondrial markers, but without genomic-level sequence data the demographic history was previously unknown.

To infer the population history and evolutionary adaptations of this endangered species, we sequenced the first red-crowned crane whole genome and compared it to other avian species. The Avian Phylogenomics Project (http://avian.genomics.cn/) previously published 48 avian reference genomes [12, 13], including the grey-crowned crane (*Balearica regulorum*) and the common ostrich (*Struthio camelus*). The grey-crowned crane belongs to the family of Gruidae together with the red-crowned crane, and is similarly proportioned (weighing up to 3.6 kg, height up to 1.4 m) [14]. However, the bodyweight of the grey-crowned crane is roughly half that of the red-crowned crane and the lifespan is considerably shorter; especially in captivity (21 years versus 65) [2]. Conversely, the common ostrich is a member of the infraclass Palaeognathae and is deeply divergent from the red-crowned crane (111 MYA). It is the largest Aves (up to 156.8 kg, up to 1.7 m in height) [15] with a lifespan closer to the red-crowned crane (maximum longevity of 50 years in captivity) [2, 15]. To identify common evolutionary signatures associated with energy metabolism and lifespan in birds, we compared our red-crowned crane genome to both the common ostrich (large/long-lived, but belonging to different family) and grey-crowned crane (belonging to the same family, but small/short-lived) genomes.

## Results

### The red-crowned crane genome

Genomic DNA from a red-crowned crane individual was sequenced using the Illumina HiSeq2000 platform (Illumina, San Diego, CA, USA). We produced a total of 494 million paired reads with a 100 bp read length and an insert size of 216 bp (Supplementary Fig. S1, Supplementary Table S2). A *K*-mer analysis (*K* = 17) estimated the genome size to be roughly 1.14 Gb (Supplementary Fig. S2, Supplementary Table S3), which is comparable to other avian species (1.0 Gb-1.2 Gb) [16]. The reads were assembled by mapping to the closely related grey-crowned crane (*Balearica regulorum*) reference genome [17]; resulting in 49 × coverage with an 81.07% mapping rate, and coding sequence (CDS) coverage of 93.48% (Supplementary Table S4). Compared to the grey-crowned crane reference genome, approximately 36 million single nucleotide variations (SNVs) and 3.6 million small insertions/deletions (indels) were identified (Supplementary Table S5). These SNVs were comprised of 34,680,112 (96%) homozygous and 1,354,842 (4%) heterozygous SNVs, yielding a nucleotide diversity of 0.00129. Compared with the median nucleotide diversity of 0.00290 in other avian species, the red-crowned crane possesses a low level of nucleotide diversity and is genetically consistent with an endangered species. A consensus sequence of the red-crowned crane was created by substituting the red-crowned crane SNVs into the grey-crowned crane reference genome (see Materials and Methods).

### Comparative evolutionary analyses of the red-crowned crane

To investigate the environmental adaptations of the red-crowned crane and the genetic basis of its longevity, we analyzed the red-crowned crane consensus sequence compared to 18 other avian genomes. We identified 7,455 orthologous gene clusters between the red-crowned crane consensus sequence and orthologous gene families from the 18 genomes (American flamingo, Anna’s hummingbird, bald eagle, budgerigar, common cuckoo, common ostrich, downy woodpecker, emperor penguin, grey-crowned crane, hoatzin, killdeer, little egret, duck, peregrine falcon, chicken, rock dove, white-tailed tropicbird, and white-throated tinamou) (Fig. 1a, Supplementary Table S6). A previously reported phylogenetic topology of these avian genomes was used in our Bayesian phylogenomic analysis [13], which estimated the divergence time between the red-crowned and grey-crowned cranes to be 20 million years ago (MYA) (Fig. 1b); in accordance with previous results [18]. The divergence time between the red-crowned crane and the common ostrich (used as an outgroup in this study) was estimated to be roughly 111 MYA.

**Figure 1.**
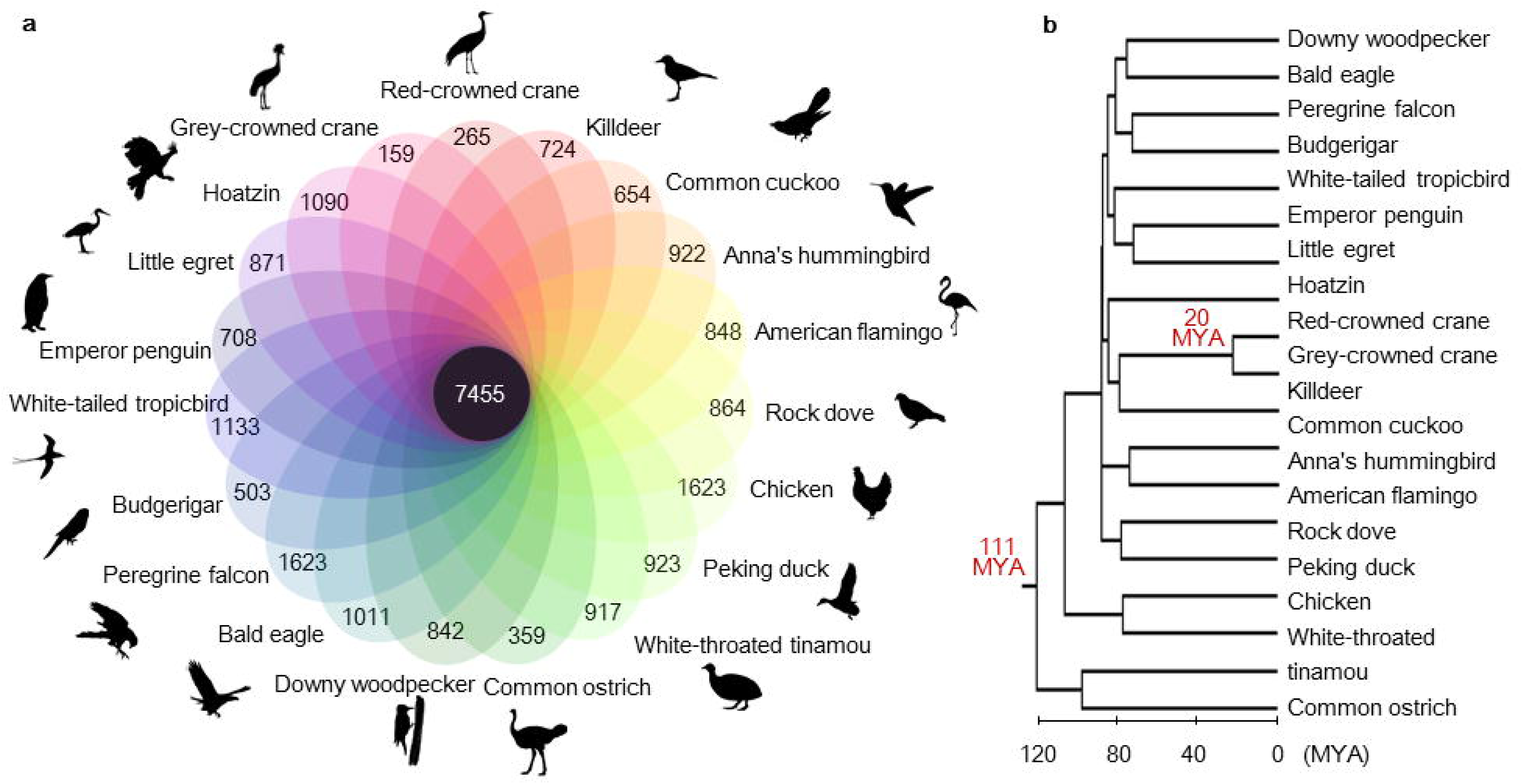
Orthologous gene clusters of the red-crowned crane compared to other avian species. (A) The Venn diagram shows the number of unique and shared gene families among the red-crowned crane and the other 18 avian genomes. (B) Divergence time of the red-crowned crane and the 18 avian species.

Avian body sizes are known to be positively correlated with lifespan and negatively correlated with metabolic rate [6, 7]. In order to investigate the contribution of genes in these pathways to the longevity of the red-crowned crane, positively selected genes (PSGs) were identified using(the branch-site and branch statistical modeling tests. We identified 3 and 162 genes as PSGs in the red-crowned crane when using a branch-site model and a branch model (1 < *d*_*N*_/*d*_*S*_ ≤5 was interpreted as positive selection), respectively (Supplementary Tables S7 and Tables S8). Of these 165 PSGs, we identified the PSGs associated with metabolism based on the KEGG pathway database to gain insight into the longevity of the red-crowned crane and the relation to metabolism (Table 1) [18]. In total, 7 PSGs *(HNMT, HS2ST1, NDUFA5, NDUFA8, NUDT12, PPCDC*, and *PSTK)* were involved in metabolic pathways. Of these, two genes were associated with energy metabolism, including *NDUFA5* ( ω average: 0.19652, ω red-crowned crane: 2.2221) and *NDUFA8* ( ω average: 0.13284, ω red-crowned crane: 1.1281). *NDUFA5* and *NDUFA8* encode the Q-module and ND1 module of NADH-ubiquinone oxidoreductase (Complex 1; EC 1.6.5.3), respectively (Fig. 2). Complex 1 functions in the transfer of electrons derived by NADH to the respiratory chain, and is a major source of damaging reactive oxygen species (ROS) that are known to be involved in aging and energy metabolism processes [19, 20]. Complex 1 is the largest protein complex in mitochondria and is perhaps the most important set of proteins in energy metabolism across all life on Earth. It is composed of N and Q modules. The N module has a NADH oxidation site with a FMN molecule as the primary electron acceptor, while the Q module has a ubiquinone reduction site. The Q-module links the matrix and mitochondrial membrane arms and is involved in transfer of electrons along Fe-S clusters to ubiquinone [19, 21]. The ND1 module is located at the base of the Q-module and is an electron transport subunit embedded in the mitochondrial inner membrane [21]. Previous studies with *NDUFA5* mutants in *Caenorhabditis elegans* have shown this gene to be is involved in lifespan extension in the nematode and the regulation of lifespan in yeast [22, 3]. Furthermore, it was reported that expression levels of both *NDUFA5* and *NDUFA8* are decreased in the mitochondria of older humans [24].

**Figure 2.**
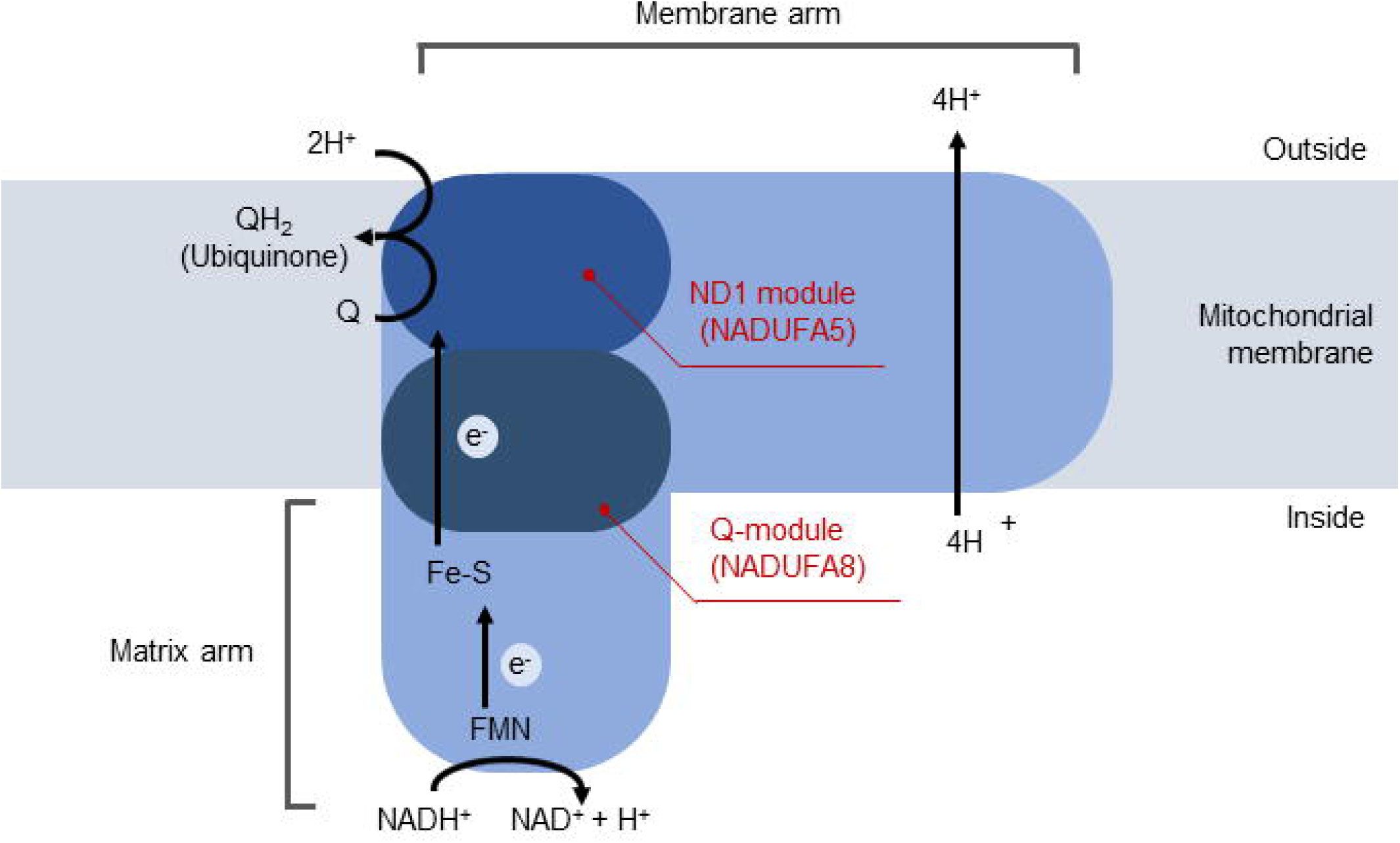
Schematic overview of the protein structure and electron transfer in NADH-ubiquinone oxidoreductase. The red line indicates the location of the ND1 module and Q-module.

**Table 1.** PSGs involved in pathways related to metabolism in the red-crowned crane.

|  |  |
| --- | --- |
| KEGG pathways | BranchModel |
| Histidine metabolism | *HNMT* |
| Oxidative phosphorylation | *NDUFA8* |
|  | *NDUFA5* |
| Glycosaminoglycan biosynthesis - heparan sulfate / heparin | *HS2ST1* |
| Metabolic pathways | *NUDT12* |
|  | *NDUFA8* |
|  | *NDUFA5* |
|  | *PPCDC* |
| Nicotinate and nicotinamide metabolism | *NUDT12* |
| Pantothenate and CoA biosynthesis | *PPCDC* |
| Selenocompound metabolism | *PSTK* |

A functional enrichment analysis of the PSGs was conducted using the DAVID functional annotation tool for Gene Ontology (GO) categories and the Fisher’s exact tests for the KEGG pathways (Supplementary Table S9 and Table S10) [18, 25]. In total, two GO terms were identified as enriched when using 10% false discovery rate. These include ribonucleoprotein complex biogenesis (GO:0022613; *EBNA1BP2*, *PHAX*, *PDCD11*, *LSM6*, and *FTSJ2*, *P* = 0.003) and ncRNA processing (GO:0034470; *PDCD11*, *LSM6*, *RG9MTD1*, *FTSJ2*, and *RPP14*, *P* = 0.004). In a previous study in humans it was discovered that the phosphorylated adapter RNA export protein encoded by the *PHAX* gene interacts with the TERT protein [26, 27, 28]. Telomere length has been proposed to be a ‘molecular clock’ that underlies organismal aging [26, 29]. The KEGG pathway enrichment test revealed five pathways with p-values < 0.05 for the PSGs determined by the branch model. They include hematopoietic cell lineage (ko04640), cell adhesion molecules (CAMs) (ko04514), primary immunodeficiency (ko05340), antigen processing and presentation (ko04612), and T cell receptor signaling pathway (ko04660). The hematopoietic cell lineage (p-value = 1.5E-05, the q-value = 6.7E-05) was the most significant KEGG pathway and includes six PSGs: *CD8B, GP9, IL9R, LOC102094583, LOC104630894*, and *LOC104643762*.

To more robustly examine the metabolic polymorphisms in this species, we identified species-specific amino acid (SSAA) changes for the red-crowned crane compared to the other 18 avian genomes. A total of 870 SSAAs were found in 696 genes (Supplementary Table S11), which were assessed for functional significance using the programs PolyPhen-2 (score > 0.15) and PROVEAN (score ≤ −2.5) [30, 31, 32]. Notably, positively selected *IL9R* gene was also found to be significantly function-altered by the SSAA analysis (polyphen-2 score = 0.858). The *IL9R* gene, which belongs to the hematopoietic cell lineage pathway, encodes a cytokine receptor that mediates the interleukin-9 response. The overexpression of *IL9R* has been shown to intensify the immune response by increasing antibody production [33]. Numerous reports indicate that immune functions are associated with aging, and most are pathologically related [34]. Among the 696 genes, 35 belonged to the KEGG metabolic pathway (Supplementary Table S12).

In addition to the evolutionary analyses for the red-crowned crane, we examined Gruidae-specific amino acid variants to gain insight into the environmental adaptations of the Gruidae family (red-crowned and grey-crowned cranes). In total, 2,416 amino acid changes were specific to the Gruidae, which were identified in 1,128 genes (Supplementary Table S13). Of these genes, only *COPS3* contained Gruidae-specific mutations (PROVEAN score = −3.131) compared to the 17 other birds (Fig. 3). COPS3 is a component of a 26S proteasome regulatory complex that has an association with muscle protein degradation [35], and previous studies have shown it to be critical for heart and skeletal muscle development in humans [36]. It is possible that this mutation is an adaptation for long body height and may play a role in providing sufficient oxygen to enable sustained powered flight [37].

**Figure 3.**
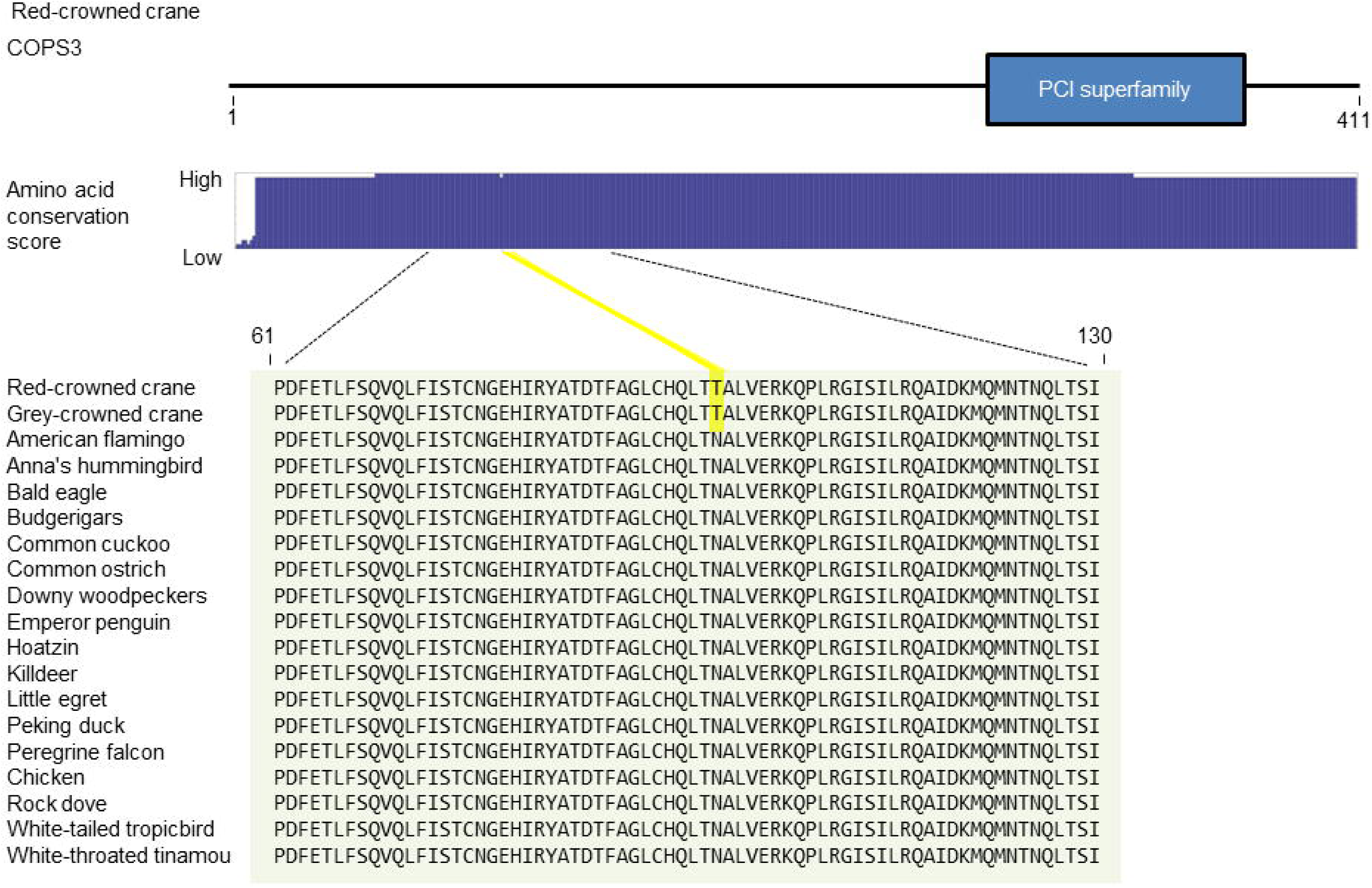
COPS3 amino acid sequence. The protein sequences from the 18 birds were aligned with that of the red-crowned crane. The yellow box represents SSAA change shared between the red-crowned crane and grey-crowned crane.

### Comparative genetic analyses of metabolic genes among long-lived Aves

The common ostrich has a lifespan comparable to the red-crowned crane [2], and we suspect that the longevity of these species could be due in part to convergent polymorphisms affecting the rate of energy metabolism. To identify any commonly shared adaptations for longevity-related physiological characteristics we compared the PSGs of the red-crowned crane with that of the common ostrich. In total, we detected five PSGs that were common between them: *NOVA1*, *SOD3*, *SOX14*, *TMEM80*, and *UBXN11* (Table 2).

**Table 2.** PSGs by branch and branch-site models that were shared in both red-crowned crane and common ostrich. M0 denotes *d*_*N*_/*d*_*S*_ for all the species in the study; M1 denotes *d*_*N*_/*d*_*S*_ for the lineage leading to the target species; Likelihood Ratio Tests (LRT) were used to detect positive selection.

| Gene | Red-crowned crane |  |  |  | Common ostrich |  |  |  |
| --- | --- | --- | --- | --- | --- | --- | --- | --- |
|  | Branch models |  | Branch-site models |  | Branch models |  | Branch-site models |  |
|  | M0:<br>one-ratio | M1:<br>free-ratio | 2ΔL | P-value | M0:<br>one-ratio | M1:<br>free-ratio | 2ΔL | P-value |
| *KIAA1009* | 0.4014 | 1.0295 |  |  |  |  | 10.8273 | 0.0155 |
| *NOVA1* | 0.1016 | 1.5792 |  |  | 0.1016 | 1.8746 | 7.9132 | 0.0489 |
| *SOD3* | 0.1454 | 1.2380 |  |  | 0.1454 | 1.4134 |  |  |
| *SOX14* | 0.0186 | 1.4472 |  |  | 0.0186 | 1.6574 |  |  |
| *TMEM80* | 0.2138 | 2.1448 |  |  | 0.2138 | 2.6535 |  |  |
| *UBXN11* | 0.4847 | 2.1552 |  |  | 0.4847 | 1.3068 |  |  |

*SOD3* is associated with the superoxide metabolic process (GO:0006801) GO molecular function [38, 39]. The free-radical theory is one of the major hypotheses of biological aging, and suggests that aging is a result of accumulated damage from free radicals of reactive oxygen [8]. ROS can be divided into two types: oxygen-centered radicals and oxygen-centered non-radicals. Oxygen-centered radicals are generally unstable and more highly reactive than oxygen-centered non-radicals. Oxygen-centered radicals include the superoxide anion radical (O_2_¯) and hydroxyl radical (OH), while oxygen-centered non-radicals include hydrogen peroxide (H_2_O_2_) and singlet oxygen (^1^O_2_) [40]. Organisms have a variety of protection mechanisms to mitigate the deleterious effects of the ROS [41]. SOD3 is an antioxidant enzyme that converts the superoxide anion radical into hydrogen peroxide (Fig. 4a) [42, 43]. In addition, the overexpression of *SOD3* results in increased cell growth and proliferation, and the inhibition of apoptosis [44]. We speculate that the *SOD3* gene product may help protect the red-crowned crane’s tissues and organs from oxidative stress, consequently increasing its longevity.

**Figure 4.**
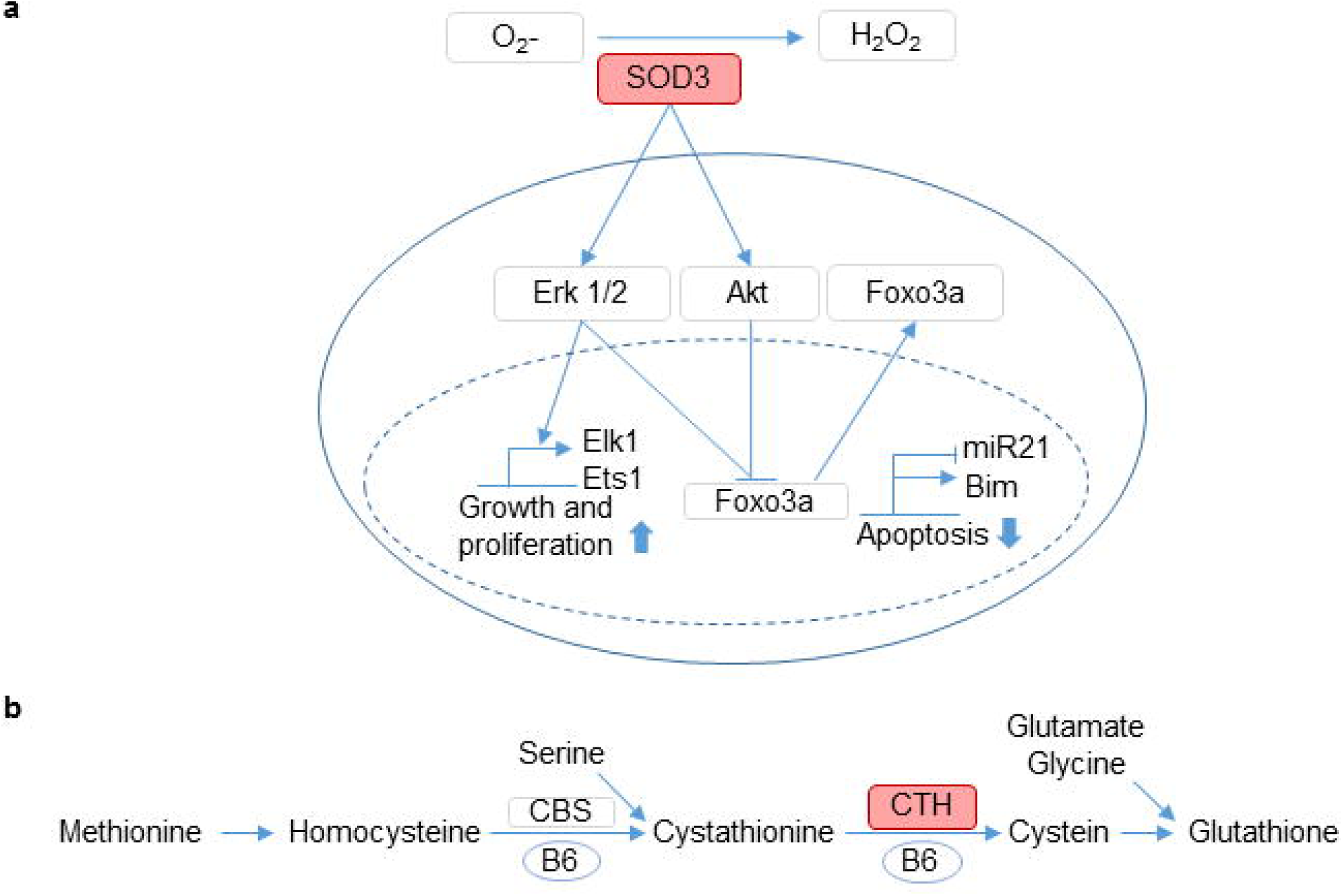
Schematic overview of signal transduction pathways. (a) Depiction of cellular responses to *SOD3*. (b) Depiction of a homocysteine metabolic pathway involving *CTH*.

In addition to comparing the red-crowned crane and common ostrich PSGs, we also identified shared amino acid changes between these two long-lived species. A total of 128 and 10,126 genes had amino acid changes specific to the red-crowned crane and the common ostrich, respectively. Among these genes, 82 were shared by the red-crowned crane and common ostrich (Supplementary Table S14). To identify candidate genes for possible convergent metabolic adaptive evolution, we predicted the functional effects of these amino acid substitutions using the programs PolyPhen-2 and PROVEAN [30, 31, 32]. We selected the genes containing deleterious variants that are possibly or probably damaging and filtered out the genes that did not belong to the KEGG metabolism pathway. In total, we discovered 17 genes that share significant amino acid changes (Table 3). Interestingly, the *NUDT12* and *PNLIP* genes showed amino acid changes at the same positions in both the red-crowned crane and common ostrich. *NUDT12* is known to regulate the concentration of the peroxisomal nucleotide cofactor for oxidative metabolism [45]. The peroxisome is clearly integrated into an endomembrane system that regulates cell senescence [46]. *PNLIP* is involved in dietary fat metabolism system and hydrolyzes dietary long-chain fatty acids to produce 2-monoacylglycerols and free fatty acids [45, 47]. Additionally, among the 18 genes with amino acid changes, *CTH* and *RPA1* genes were listed in the GenAge database as aging-related genes [48]. *CTH* encodes a cytoplasmic enzyme for the methionine pathway (Fig. 4b). Methionine is an essential amino acid found mainly in animal protein. Homocysteine is converted from methionine, which has a direct toxic effect on endothelial cells, and increased homocysteine synthesis is associated with aging [49, 50]. *CTH* is required for the conversion of methionine into cysteine, which is necessary for the synthesis of the antioxidant glutathione [51]. In addition, it was reported that RNAi-mediated knockdown of *CTH-1* and *CTH-2*, which are known as pro-longevity genes, in *C. elegans* caused median lifespan reductions (up to ~15%) [52]. *RPA1* encodes an RPA heterotrimeric complex and three functional domains [53] which interacts with aging-related genes such as *WRN* [54]. The amino acid substitution sites are included in a ssDNA-binding domain that is involved DNA replication, recombination, and repair [55].

**Table 3.** Unique amino acid changes of the red-crowned crane and common ostrich in metabolism related proteins.

| Protein | Red-crowned crane |  |  | Common ostrich |  |  |
| --- | --- | --- | --- | --- | --- | --- |
|  | Position | PROVEAN (score) | Polyphen (score) | Position | PROVEAN (score) | Polyphen (score) |
| AGL | Y1067C | Deleterious (-3.446) | Benign (0) | V690A | Deleterious (-3.367) | Benign (0.001) |
|  |  |  |  | C1164S | Deleterious (-5.536) | Benign (0) |
|  |  |  |  | P1277A | Deleterious (-7.107) | Benign (0.001) |
|  |  |  |  | R1486H | Deleterious (-2.864) | Benign (0.001) |
| AK9 | P366S | Deleterious (-7.884) | possibly damaging (0.599) | E310A | Deleterious (-5.783) | Benign (0.025) |
|  |  |  |  | K271R | Deleterious (-1.721) | Benign (0.001) |
| ALG1 | R93H | Deleterious (-3.687) | Benign (0.102) | L99F | Deleterious (-2.869) | Benign (0.024) |
|  | I298M | Neutral (-2.322) | possibly damaging (0.857) |  |  |  |
| CTH | T366A | Deleterious (-4.058) | Benign (0.095) | A140S | Deleterious (-2.718) | Benign (0.031) |
| EPRS | K1068R | Neutral (-1.497) | possibly damaging (0.872) | T613C | Deleterious (-4.198) | Benign (0.013) |
| FMN1 | Q40E | Deleterious (-2.773) | possibly damaging (0.659) | N14S | Deleterious (-3.917) | Benign (0.003) |
| GNE | L501I | Neutral (-0.492) | probably damaging (0.98) | I216M | Neutral (-0.657) | possibly damaging (0.861) |
| HAAO | F186V | Deleterious (-3.216) | probably damaging (1) | L170P | Deleterious (-3.348) | Benign (0.005) |
| HIBCH | T119M | Deleterious (-3.750) | probably damaging (0.989) | S223F | Deleterious (-4.536) | Benign (0.009) |
| MOCS2 | S187L | Deleterious (-3.219) | Benign (0.115) | V151T | Deleterious (-3.814) | Benign (0.013) |
| NUDT12 | Q170P | Deleterious (-2.811) | Benign (0.003) | E297G | Deleterious (-5.273) | Benign (0.016) |
|  | E364K | Deleterious (-3.993) | probably damaging (0.988) |  |  |  |
| PHOSPHO2 | Y133C | Deleterious (-4.009) | Benign (0.007) | S38C | Deleterious (-3.320) | Benign (0.255) |
|  |  |  |  | R50K | Deleterious (-2.714) | Benign (0.136) |
|  |  |  |  | V156A | Deleterious (-3.487) | Benign (0.064) |
| PIGL | L49V | Neutral (-2.080) | probably damaging (1) | P127S | Deleterious (-2.080) | Benign (0.023) |
| PNLIP | E17G | Deleterious (-5.347) | Benign (0.054) | E17G | Deleterious (-5.347) | Benign (0.054) |
|  | Y105H | Neutral (0.958) | probably damaging (0.988) |  |  |  |
| QRSL1 | V97A | Deleterious (-3.633) | probably damaging (0.99) | H341Y | Deleterious (-3.186) | Benign (0.012) |
| RPA1 | T197M | Deleterious (-4.899) | probably damaging (1) | P127A | Deleterious (-3.928) | Benign (0) |
|  |  |  |  | N495T | Deleterious (-5.190) | Benign (0) |
| UMPS | Q417E | Deleterious (-2.683) | possibly damaging (0.539) | E193G | Deleterious (-2.994) | Benign (0.002) |

### Genetic diversity and population history

The pairwise sequentially Markovian coalescent (PSMC) analysis was used to model the historical population size fluctuations of the red-crowned crane and the closely related grey-crowned crane (Fig. 5) [56]. We discovered a striking difference in effective population size between these two species during last glacial age. While the effective population size of the red-crowned crane appears to have peaked about 80,000 years ago in the last glacial age, the effective population size of the grey-crowned crane was expanded to their maximum roughly 500,000 years ago.

**Figure 5.**
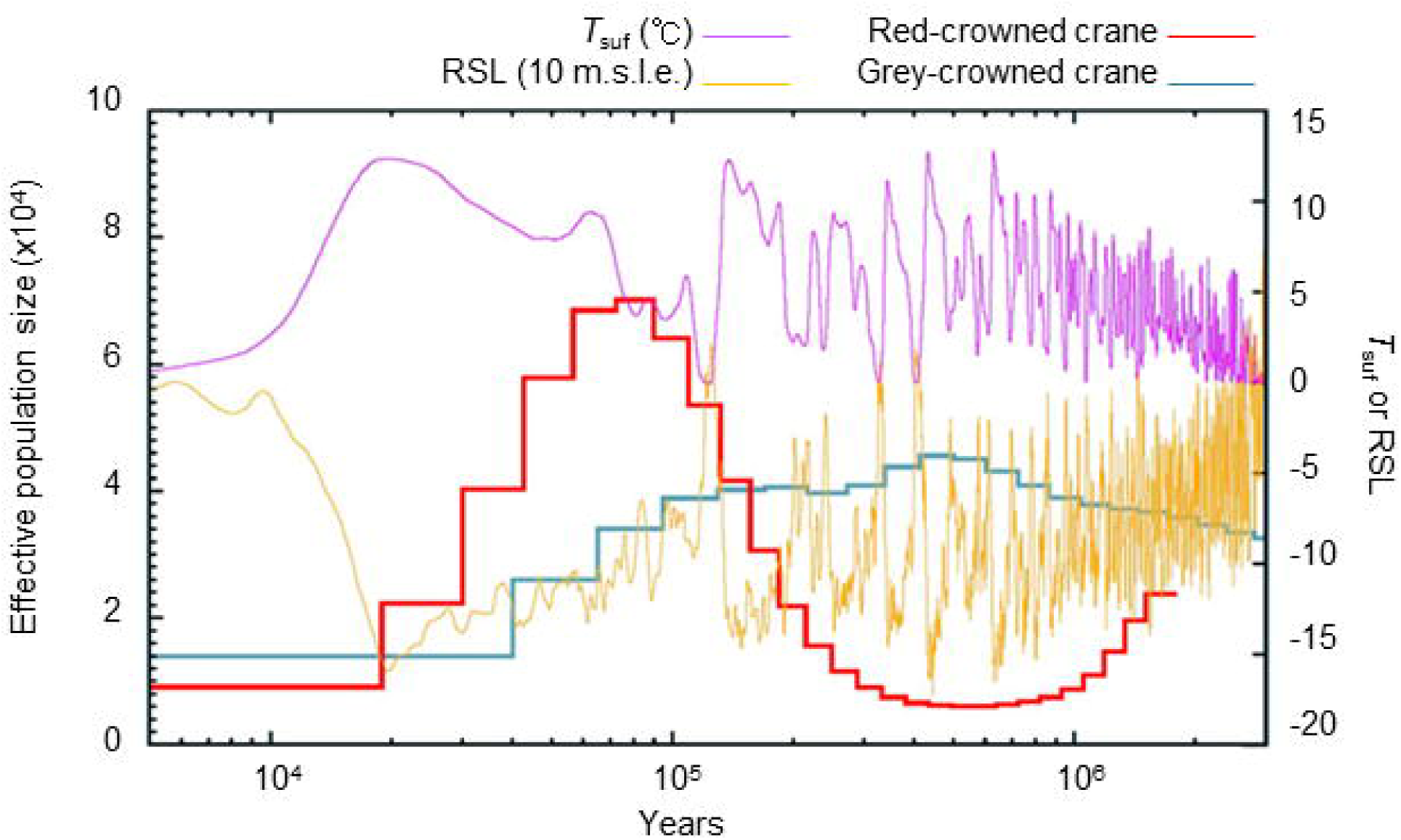
Demographic history of the red-crowned crane and grey-crowned crane. Red-crowned crane PSMC model with the generation time (g=12.3) and mutation rate (μ= 9.66 × 10^−9^), and grey-crowned crane PSMC model with the generation time (g= 15.1) and mutation rate (μ= 1.18 × 10^−8^). *T*_*suf*_, atmospheric surface air temperature; RSL, relative sea level; 10 m.s.l.e., 10 m sea level equivalent.

## Discussion

We generated and analyzed the first whole genome sequence of the red-crowned crane. By comparing the red-crowned crane whole genome sequences to the grey-crowned crane reference and other avian genomes, we identified *NDUFA5* and *NDUFA8* genes as positively selected in the red-crowned crane, which are associated with the mitochondrial respiratory chain complex 1 involved in energy metabolism and longevity. In addition, the *IL9R* gene, which is associated with immunity, was identified as both a PSG and a gene that contains red-crowned crane-specific amino acid changes, which may play a role in its longevity. We also identified a significant SSAA change in the COPS3 protein that is involved in cardiac and skeletal muscle development, which may be an adaptation for long body height the family Gruidae. We suspect that, as in the giraffe, cardiovascular muscle associated gene like COPS3 help maintain blood delivery to the brain despite their long and narrow necks. Taken together, we believe that these candidate genes warrant further experimental study to test their impact on longevity and physiological effects.

We identified common signatures of longevity-related adaptive evolution between the red-crowned crane and common ostrich and identified shared positively selected genes that significantly associated with metabolism and lifespan. Of these candidate genes, *SOD3* is involved in the superoxide metabolic process. Additionally, the *NUDT12* and *PNLIP* genes showed significant SSAA changes at the same position in both the red-crowned crane and common ostrich. *NUDT12* is associated with the metabolism of peroxisome, which might contribute indirectly to aging. Furthermore, SSAA changes of *CTH* and *RPA1* are also associated with longevity-related metabolic pathways. Of special interest are the amino acid substitutions at identical loci in *NUDT12* and *PNLIP*, which may provide additional examples of convergent adaptive evolution through parallel amino acid replacements. Similar signatures have been observed, and then experimentally verified, in the hemoglobin sequences of some high-altitude birds [57]

Demographic modeling showed a population decline of the red-crowned crane since the last glacial period roughly 80,000 years ago, and we calculated a low nucleotide diversity of 0.00129. While it is possible that their northern habitat naturally suppresses their effective population size, it is also known that habitat loss and pollution have contributed to severe population declines. Taken together, these findings further highlight the importance of continued monitoring and management of this endangered species. We hope the consensus genome presented here will be a valuable resource for future conservation genetic studies of this iconic species.

## Methods

### Sample preparation, genomic DNA extraction, and sequencing

The protocol of the blood sample preparation was carried out in accordance with guidelines of Korean Association for Bird Protection under the Cultural Heritage Administration (Korea) permit. All experimental protocols were approved by the Genome Research Foundation. A blood sample was secured from a single female red-crowned crane collected upstream of Gunnam dam, Gyeonggi-do, Republic of Korea (Lat = 38°06′24.8″N and Long = 127°01′17.1″E). Genomic DNA was extracted using a QIAamp DNA Mini Kit (Qiagen, CA, USA) following the manufacturer’s instructions. The DNA concentration was measured using the Qubit dsDNA assay kit (Invitrogen, CA, USA) and an Infinite 200 PRO Nanoquant system (TECAN, Mannedorf, Germany). Fragmentation of high-molecular weight genomic DNA was carried out with a Covaris S2 Ultrasonicator (Covaris, Woburn, MA, USA), generating 500 bp fragments. Whole-genome shotgun libraries were prepared according to Illumina’s library preparation protocols using a TruSeq library sample prep kit (Illumina, CA, USA). Aliquots were analyzed on an Agilent 2100 Bioanalyzer (Agilent Technologies, CA, USA) to determine the library concentration and size. Sequencing was performed on an Illumina HiSeq 2000 sequencer (Illumina, CA, USA), using the TruSeq Paired-End Cluster Kit v3 (Illumina, CA, USA) and the TruSeq SBS HS Kit v3 (Illumina, CA, USA) for 200 cycles.

### Red-crowned crane genome assembly

Low quality DNA reads with phred quality scores <20 and/or ambiguous nucleotides ‘N’ ratios >10% were filtered out. Clean reads were aligned to the grey-crowned crane genome sequence using BWA 0.6.2 [58] at default settings. PCR duplicates from the reads were removed using ‘rmdup’ command of SAMtools 0.1.18 [59] at default settings. Complete consensus sequences were determined by SAMtools mpileup, Bcftools view, and SAMtools vcfutils.pl vcf2fq pipelines from the SAMtools 0.1.19 suite [59]. The consensus sequence included all alignment depths. SNV calling was conducted based on the consensus genome sequences. Indels were called using the GATK 3.3 UnifiedGenotyper with-dcov 10000 option. The detected indels were marked by VariantFilteration in GATK 3.3 with the following criteria: (1) Hard to validate, MQ0≥4 && ((MQ0/ (1.0 * DP)) >0.1); (2) Quality filter, QUAL <10.0; (3) Depth filter, DP < 5. The SnpEff 3.3 software was used to predict the effects of the indels. Heterozygous SNV rate was used as the nucleotide diversity value. Other avian nucleotide diversity was calculated as an average value of the American flamingo (0.00372), Anna’s hummingbird (0.00288), common ostrich (0.00176), downy woodpecker (0.00455), grey-crowned crane (0.00202), peregrine falcon (0.00112), white-throated tinamou (0.00560) and White-tailed tropicbird (0.00162). CDS were selected by filtering out the following conditions: (1) there is an ambiguous nucleotide ‘N’ in a CDS; (2) there is a premature stop codon in a CDS. Among the 14,173 CDSs, 13,407 CDSs were selected and used in subsequent analyses. *K*-mers (*K* = 17) were obtained using the JELLYFISH 2.2.4 [60] program with the command “jellyfish count-m 17-o output-c 32-s 8589934592-t 64”. We calculated the genome size from the total *K*-mers frequencies.

### Orthologous gene clustering

The protein sequences of the red-crowned crane were predicted using the gene model of the grey-crowned crane. Reference genomes of the grey-crowned crane and more distant 18 avian genomes were downloaded from the GigaDB dataset (http://gigadb.org/dataset/101000). The clusters of orthologous genes in the 19 genomes were constructed using OrthoMCL 2.0.9 [61]. Self-to-self BLASTP searches were conducted for all the protein sequences of the red-crowned crane and 18 avian genomes with an E-value of 1E-5. The Markov clustering algorithm was used to define the cluster structure between the proteins with inflation values (-I) of 1.5. Lastly, the OrthoMCL program was used to determine orthologous gene clusters with an e-value exponent cutoff value of −5 and percent match value of 50%. In total, we identified 7,455 one-to-one orthologous groups from the 19 avian genomes. Figure 1 and 2 were drawn manually using the online photo editor (https://pixlr.com/editor/).

### Evolutionary analyses

We estimated divergence times for the red-crowned crane and 18 avian reference genomes. The white-throated tinamou’s divergence time was taken from TimeTree [17], and the others were calculated using MEGA7 RelTime-CC 2.0 program with the topology from the phylogenetic tree published previously [62]. The reference data for this analysis included the common ostrich (111 MYA), grey-crowned crane (98 MYA) and emperor penguin (81 MYA), compared to the red-crowned crane. The maximum likelihood statistical method was used based on a general time reversible model. The multiple sequence alignment of orthologous genes was constructed using PRANK alignment program package [63], and the rates of synonymous (*d*_*S*_) and non-synonymous substitutions (*d*_*N*_) and the *d*_*N*_/*d*_*S*_ ratio (nonsynonymous substitutions per nonsynonymous site to synonymous substitutions per synonymous site, ω) were estimated using the CODEML program in PAML 4.5 [64]. The one-ratio model (null model) detects the single *d*_*N*_/*d*_*S*_ ratio for all the branches of a phylogenetic tree. It was used to measure the general selective pressure acting on all of the species. A free-ratios model assumes the *d*_*N*_/*d*_*S*_ ratio along each branch. The branch-site test of positive selection based on the likelihood ratio test (LRT) was conducted with a conservative 10% false discovery rate criterion. The Gruidae and common ostrich were used as the foreground branch for each analysis accordingly, and the other species were used as background branches. Functional effects of the SSAAs changes were predicted using PolyPhen-2 and PROVEAN [30, 31, 32] programs using the default cutoff values.

### Demographic history

Whole genome sequencing data can be used to accurately estimate the population history [65, 66, 67]. We inferred the demographic history of the red-crowned crane using a PSMC analysis with scaffolds of length ≥50 kb. The PSMC program was performed with 100 bootstrapping rounds, and run with options of 9.59 × 10^−9^ substitutions per site per generation [12] and generation time of 12.3 years as previously reported [9].

## Availability of supporting data

Whole-genome sequence data was deposited in the SRA database at NCBI with BioSample accession number SAMN07580860. The whole genome sequencing can be accessed via reference number SRX3148473. The data can also be accessed through BioProject accession number PRJNA400839 for the whole-genome sequence data.

## Acknowledgments

This study was supported by PGI of Genome Research Foundation and Geromics Inc. internal research funds and the No.10075262 of National Center for standard Reference Data. This research was also supported by Ulsan National Institute of Science & Technology (UNIST) internal research funds. And the National Research Foundation of Korea (2013M3A9A5047052 and 2017M3A9A5048999) research funds. The authors thank many people not listed as authors who provided analyses, data, feedback, samples, and encouragement. Especially, thanks for Hak-Min Kim, Taehyung Kim and Byungchul Kim.

## Author contributions statement

J.B. and W.P. conceived the experiments; W.P., J.L. prepared the sample; H.L, S.J., O.C. and J.J. analyzed the data; H.L. and S.J. visualization and validation the data; H. L., S.J., O.J., J.J., Y.J. and J.B. interpreted the result; H. L. and S.J. wrote the original manuscript preparation; H.L., S.J., O.C., Y.C., J.J., J.K., J.W., J.E. and J.B. edited, and revised the manuscript; All authors read and approved the final manuscript.

## Supplementary information

1. Supplementary Information

Supplementary Table

2. Supplementary Information

Supplementary Figure

